# Introducing furanocoumarin biosynthetic genes in tomato results in coumarins accumulation and impacts growth

**DOI:** 10.1101/2025.07.07.663522

**Authors:** Alexandre Bouillé, Cloé Villard, Gianni Galati, Marwa Roumani, Aude Fauvet, Jérémy Grosjean, Lennart Hoengenaert, Wout Boerjan, John Ralph, Frédérique Hilliou, Christophe Robin, Alain Hehn, Romain Larbat

## Abstract

Over the past three decades, efforts to decipher plant metabolism have shed light on key enzymes driving specialized metabolite biosynthesis. Although only few pathways have been completely investigated to date, their characterization paves the way for exploring the potential effects of specialized metabolites on plant physiology. Among them is the linear furanocoumarin pathway, which was recently completed to produce up to psoralen. In this study, we report the first metabolic engineering of the linear furanocoumarin pathway to enable artificial psoralen production in tomato, through the integration of four genes coding for the enzymes: Umbelliferone Synthase, Demethylsuberosin Synthase, Marmesin Synthase and Psoralen Synthase. Interestingly, coumarins were produced instead of furanocoumarins. Using morphophysiological, metabolomic, and transcriptomic analyses, we suggest how coumarins, particularly scopoletin, can impact growth and affect plant physiology, even at low concentrations. As coumarins have increasingly attracted interest for agricultural applications due to their minimal environmental impact, this work both expands and challenges their potential by highlighting the physiological costs and benefits they may impose on tomato.

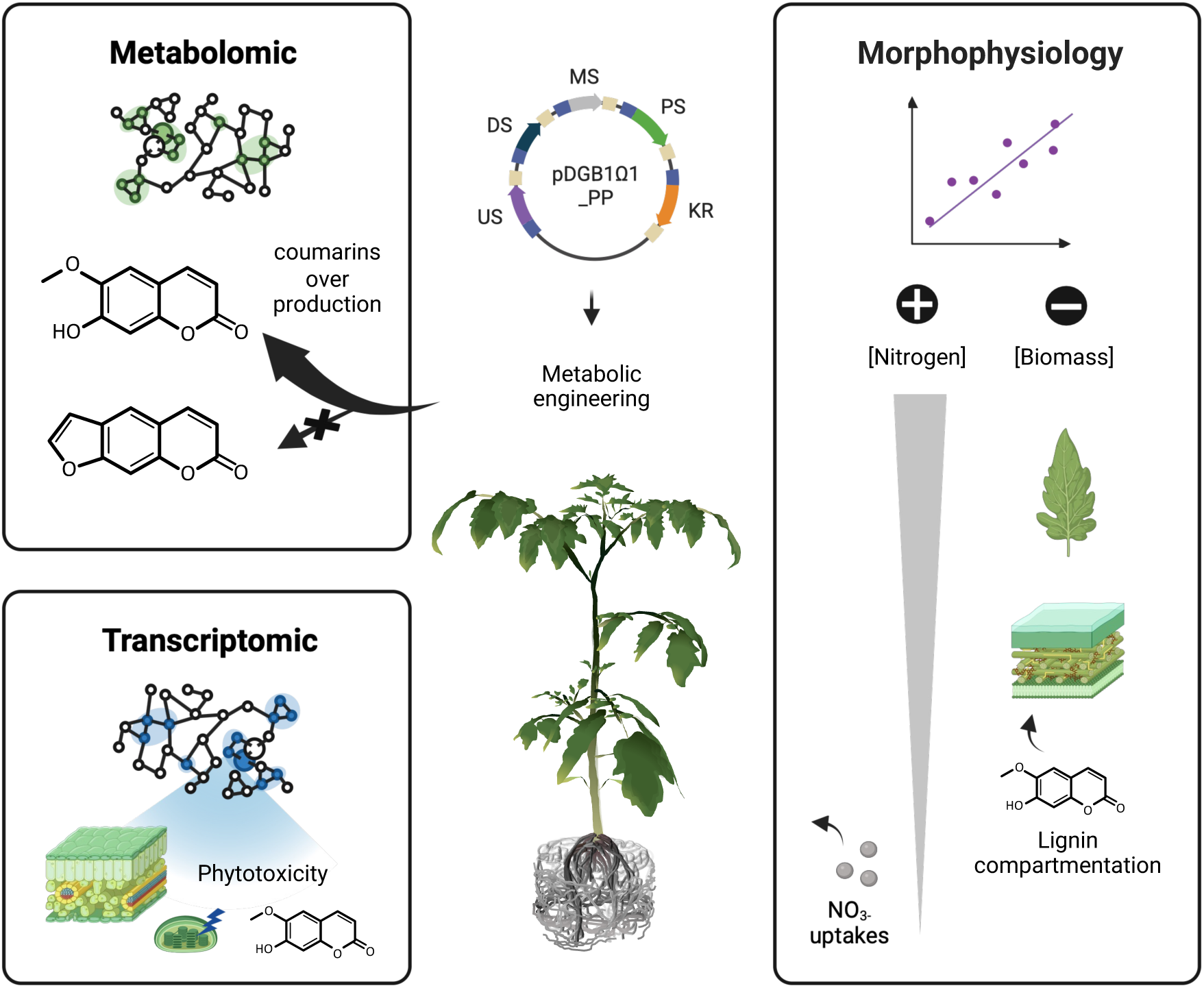
Graphical abstract.

## 1. Introduction

The around 400 million years of evolution since the terrestrialization of plants have driven the emergence of numerous mechanisms for interacting with an ever-evolving environment (Morris *et al.,* 2018). The production of specialized metabolites is a key example of such adaptation as many of these compound play vital roles in enabling plants to cope with biotic and abiotic stresses (Maeda & Fernie, 2021; Weng *et al.,* 2021). Investment of resources and energy in processes that promote plant defense, particularly the biosynthesis of specialized metabolites, may come at the expense of processes that support plant growth (He *et al.,* 2022). Plants must therefore balance the trade-oe between investing in the costly biosynthesis of specialized metabolites and allocating resources for growth (Neilson *et al.,* 2013).

Land plants produce a wide range of specialized metabolites (Chassagne *et al.,* 2019). Among them, linear furanocoumarins are defensive metabolites restricted to certain lineages (Sarker & Nahar, 2017). Its pathway has been largely elucidated over the past two decades, particularly in Apiaceae and Moraceae (**Fig. 1**) (Bouillé et al., 2025). Furanocoumarins are derived from the ubiquitous coumarin umbelliferone, which itself is derived from *p*-coumaroyl-CoA (Vialart *et al.,* 2012; Roselli *et al.,* 2017). The biosynthesis of linear furanocoumarins then starts with the C6-prenylation of umbelliferone to produce demethylsuberosin by a prenyltransferase (Karamat *et al.,* 2014; Munakata *et al.,* 2016). Demethylsuberosin is then cyclized by marmesin synthase, a cytochrome P450 (P450), into marmesin (Villard *et al.,* 2021; K. Wang *et al.,* 2024), which is subsequently converted to psoralen via C7-decarboxylation through *syn*-elimination of acetone by psoralen synthase, another P450 (Jian *et al.,* 2020; Larbat *et al.,* 2007, 2009). The psoralen backbone can then be hydroxylated at C5 or C8 by additional P450s to yield bergaptol or xanthotoxol, respectively (**Fig. 1**) (Krieger *et al.,* 2018; Ji *et al.,* 2024). These structures can be further decorated by other enzymes, such as *O*-methyltransferases, *O*-prenyltransferases and/or other P450s, to generate more complex alkylated structures (Hehmann *et al.,* 2004; Zhao *et al.,* 2016; Han *et al.,* 2023; Bouillé *et al.,* 2025). Along with some of its derivatives, psoralen exhibits a broad spectrum of defensive effects, mainly driven by its ability to form covalent bonds with nucleic acids (*i.e.,* via photocycloaddition) and to alter herbivore defences (*e.g.,* through cytochrome P450 inhibition) (Neal & Wu, 1994; Kitamura *et al.,* 2005). For such reasons, psoralen is considered the first toxic metabolite in the linear furanocoumarin pathway.

**Figure 1:**
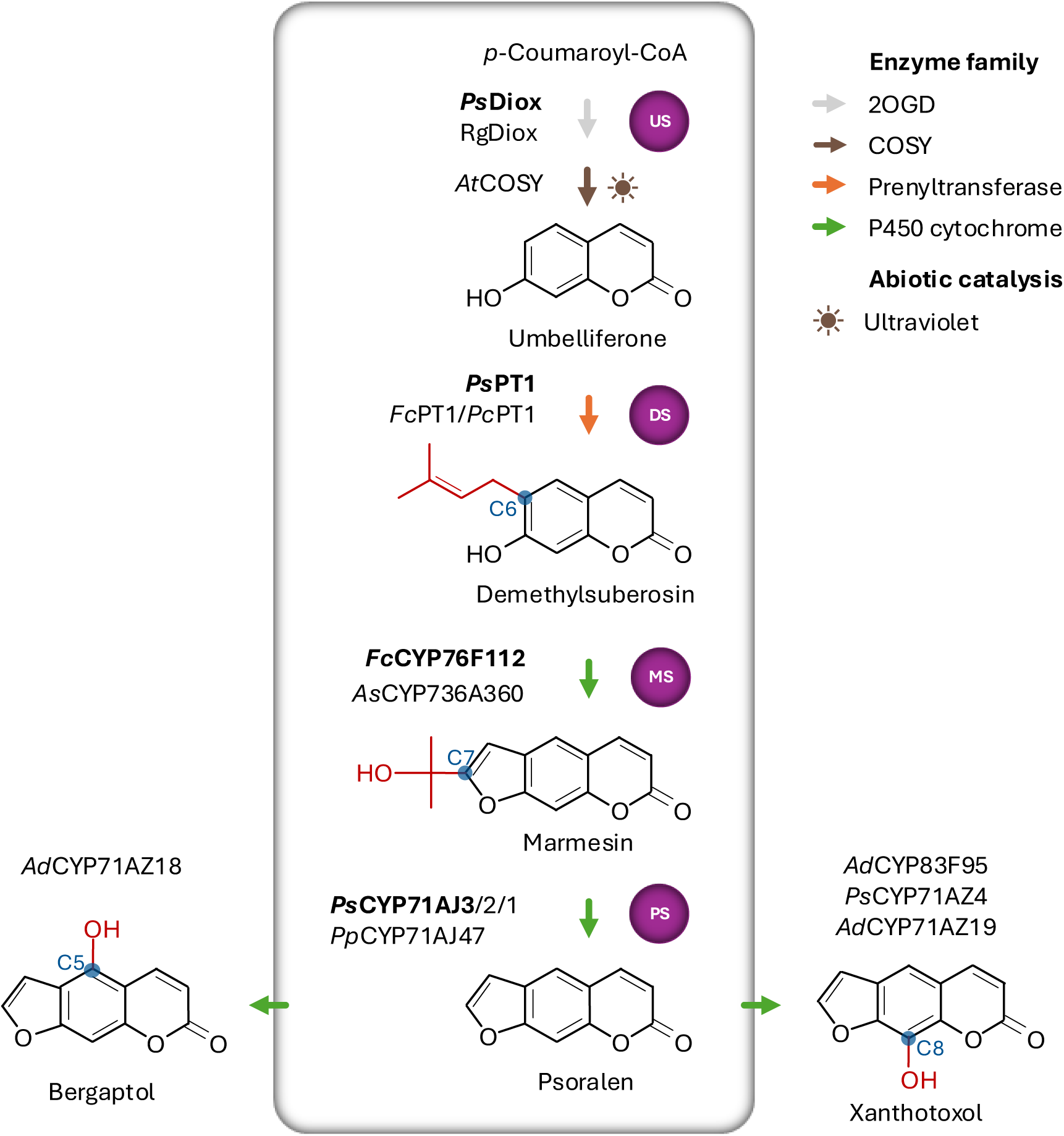
Simplified diagram of the psoralen pathway (PP) among linear furanocoumarin in some restricted angiosperms taxa. The pathway from *p*-coumaroyl-CoA to psoralen is framed, with the enzymes engineered in the tomato PP line encircled in purple and the specific genes inserted in the tomato genome in bold, while the carbon number on the furanocoumarin backbone is depicted in blue. Abbreviations: 2-OxoGlutarate dependent Dioxygenase (2OGD), COumarin SYnthase (COSY), PrenylTransferase (PT), CYtochrome P450s (CYP), US (*UMBELLIFERONE SYNTHASE*), DS (*DEMETHYLSUBEROSIN SYNTHASE*), MS (*MARMESIN SYNTHASE*), PS (*PSORALEN SYNTHASE*). Species: Apiaceae (*Angelica dahurica, Angelica sinensis*, *Pastinaca sativa*, *Petrolinum crispum*, *Peucedanum praeruptorum*), Rutaceae (*Ruta graveolens*), Brassicaceae (*Arabidopsis thaliana*), Moraceae (*Ficus carica*). From Larbat *et al.,* 2007, 2009; Vialart *et al.,* 2012; Karamat *et al.,* 2014; Munakata *et al.,* 2016; Roselli *et al.,* 2017; Vanholme *et al.,* 2019; Jian *et al.,* 2020; Villard *et al.,* 2021; Wang *et al.,* 2024.

In this study, we aimed to investigate the potential physiological and growth-related costs associated with the production of a novel specialized metabolite in plants using metabolic engineering. Considering the importance of psoralen in plant-environment interactions and the unknown costs its biosynthesis may impose on plants, we chosen to investigate the recently completed psoralen pathway. In that regard, using the GoldenBraid multigenic assembly system (Sarrion-Perdigones *et al.,* 2013), we integrated four linear furanocoumarin biosynthetic genes into the tomato genome (*Solanum lycopersicum*, Solanaceae), a plant that does not naturally produce these compounds, to enable the conversion of the ubiquitous *p*-coumaroyl-CoA into psoralen (**Fig. 1**). Metabolic analyses of the transgenic tomato line (referred here to as “Psoralen Pathway” or PP line) confirmed the detection of small quantities of psoralen, the expected product, but also highlighted larger modifications in the accumulation of other metabolite families, among them coumarins and particularly scopoletin. This modulation of the metabolic composition among PP line was accompanied by major transcriptomic changes and a significant modification in the biomass distribution. Our results shed light on the physiological costs associated with the implementation of the psoralen pathway in tomato and offer novel perspectives on the use of coumarins in agriculture.

## 2. Material and methods

### 2.1. Biological material and Standards

The tomato plants (*Solanum lycopersicum*) cultivar *West Virginia* 106 (Wva106) was used as the wild-type control and for plant transformation. Analytical standards were sourced from Sigma-Aldrich (Saint-Louis, MO, USA; for *p*-coumarate, scopoletin, and esculin), Extrasynthese (Lyon, France; for umbelliferone, psoralen, and esculetin), TransMIT (Giessen, Germany; for marmesin), Phytolab (Nantes, France; for scopolin), and R. Munakata from Kyoto University (demethylsuberosin).

### 2.2. Creation of the PP transgene

The Psoralen Pathway (PP) T-DNA was created by combining the genes controlling the 4-steps metabolic pathway from *p*-coumaroyl-CoA to psoralen (**Fig. 1**). The four genes were *Ps*Diox (ncbi: KY081678; Roselli *et al.,* 2017) coding for Umbelliferone Synthase (US), *Ps*PT1 (ncbi: KM017083; Munakata *et al.,* 2016) coding for Demethylsuberosin Synthase (DS), *Fc*CYP76F112 (ncbi: MW348922; Villard *et al.,* 2021) coding for Marmesin Synthase (MS) and *Ps*CYP71AJ3 (ncbi: C0SJS2; Larbat *et al.,* 2009) coding for Psoralen Synthase (PS). The selection marker used in the creation of the PP transgene was the *NEOMYCIN PHOSPHOTRANSFERASE* (*Ec*NPTII) gene conferring resistance to kanamycin, which originates from the plCSL11024 vector. The coding sequence of the five genes of interest were domesticated by removing internal restriction sites for *Bsa*I and *Bsm*BI, and by adding flanking extensions compatible with the GoldenBraid 2.0 multigene cloning system (GB2.0; Sarrion-Perdigones *et al.,* 2013). Each domesticated gene was cloned into the pUPD vector using the *Bsm*BI restriction enzyme, and assembled with the *Ca*MV P35S promoter (pUPD-35S, Addgene plasmid #68163) and the tNOS terminator (pUPD-tNOS, Addgene plasmid #68188) to form individual transcription unit (TU). TUs were assembled into the plasmid pDBGB1α1 or pDBGB1α2 by restriction/ligation using the restriction enzymes *Bsa*I or *Bsm*BI, respectively, and the T4 DNA ligase, as described in Sarrion-Perdigones *et al.,* 2011. The resulting TUs were further assembled two-by-two into pDGB1 α-or Ω-level plasmids, through restriction/ligation, to build the final PP T-DNA into pDGB1Ω1. Integrity of this final vector pDGB1Ω1_PP was validated by PCR analysis (see primers section 2.3), restriction analysis (**Fig. S1a**), and sequencing of the coding genes (Eco-Seq, Macrogen). PCR amplification was performed using the SapphireAmp® polymerase (Takara Bio) and following the reaction cycle: Initial denaturation (98 °C for 5 min), 35 cycles composed of a denaturation step (98 °C for 10s), an annealing step (55 °C for 15 s) and an elongation step (72 °C for 10 s), followed by a final elongation step (72 °C for 1 min). Restriction of 1μg of plasmid DNA was performed using *Bsa*I, *Bsm*BI, *Pvu*II, and *Eco*RI enzymes with the FastDigest system (Thermo Fisher Scientific), following the manufacturer’s recommendations.

### 2.3. Plant transformation and regeneration of the PP line

The pDGB1Ω1_PP vector was co-transformed into *Agrobacterium tumefaciens* EHA105 with the replication helper vector pSOUP. The transformed cells were selected on solid LB medium containing spectinomycin (100 mg/L), tetracycline (5 mg/L), and rifampicin (20mg/L) (Hellens *et al.,* 2000; Sarrion-Perdigones *et al.,* 2013). A pre-culture was performed overnight at 28 °C, 200 rpm, in 10 mL of LB medium with appropriate antibiotics. A 50 mL culture was then started with 5 mL of the pre-culture to reach an OD_600nm_ of 0.4-0.6. The culture was centrifuged for 10 min at 7000 ×g, then resuspended in 40mL of sterile water and diluted to 30mL to achieve an OD_600nm_ of 0.08. Axenic cotyledon fragments approximately 4mm in size were harvested from 7-day-old tomato seedlings grown under controlled conditions (24 °C, 16 h/8 h light/dark cycles; 150 μmol.m^-2^.s^-1^, Fitoclima 1200, Aralab). The cotyledon fragments were cultured for 24 h at 24 °C in darkness before being co-incubated with the Agrobacteria solution for 30 min at room temperature (gentle agitation every 5min). The fragments were then dried on axenic paper and placed adaxial side up on co-cultivation medium for 48 h at 24 °C in darkness. Partially transformed tissues were selected on kanamycin-enriched callus medium (150 mg/L) in Petri dishes (15 fragments per dish) at 24 °C (16 h/8 h light/dark cycles; 150 μmol.m^-2^.s^-1^) and subcultured weekly. After 3 months of subcultures, differentiated seedlings from calli were cut and transferred to rooting medium before being acclimated in soil in phytotrons (24 °C; 16 h/8 h light/dark cycles; 400 μmol.m^-2^.s^-1^; 70% humidity). Several heterozygous descendants from the regenerated tomato line were obtained by self-pollination. The insertion of the 5 transgenes into the genome was assessed by PCR (**Fig. S1b**) using genomic DNA extracted with the kit following the supplier’s recommendations and the primers described as follow: *Ps*Diox_F: CTAGTCGAAATGGCTCCATCTCC; *Ps*Diox_R: CGAAGATCCATTCCAAGCTCACATTTG; *Ps*PT1_F: CTAGTCGAAATGGCTCAAACAATTATGC; *Ps*PT1_R: GAAGATCCATTCCAAGCTCACCTC; *Fc*CYP76F112_F: CTATTCTAGTCGAAATGGATATTTTCACCTC; *Fc*CYP76F112_R: CGAAGATCCATTCCAAGCCTAATGATG; *Ps*CYP71AJ3_F: CTATTCTAGTCGAAATGAAGATGCTTGAG; *Ps*CYP71AJ3_R: GATCGAAGATCCATTCCAAGCTCATC; NPTII_F: CAATTTACTATTCTAGTCGAAATGGTTGAACAAG; NPTII_R: GAAGATCCATTCCAAGCTCAGAAGAAC. The expression of the 5 genes in each individual was confirmed by RT-PCR for each tomato line (**Fig. S1c**). Total RNAs was extracted using the E.Z.N.A.® HP kit (Omega Bio-Tek) instructions, treated with DNase (AMPD1-1KT, Sigma) to remove genomic DNA contaminants, and then reverse-transcribed into cDNA using the High-Capacity RNA-to-cDNA Kit (Applied Biosystems^TM^), according to the supplier’s recommendation.

### 2.4. Plant cultivation and measurement of morpho-physiological traits

Four sets of plant cultivation and harvest were performed over a period of four months. Three sets of 6-8 plants per genotype were used for transcriptomic and metabolomic analyses (23 WT, 19 PP), and one additional set (23 WT, 19 PP) for morphophysiological measurements, for a total of 46 WT and 38 PP plants. All plants were staggered under identical conditions. The first three were used for the metabolomics and transcriptomics analyses. The fourth series was dedicated to measuring morphometric traits, including carbon and nitrogen allocation, and lignin composition/content in the stems. At harvest, stem and petiole tissues were separated from the leaves and roots. For metabolomics analyses, petioles and internodes were combined.

Tomato plants were grown in 1.5 L pots with 0.5 L of potting soil (Stender D400, NPK 14/10/18, pH 6.2, sieved to 5 mm). The climatic conditions in the growth chamber were set at 24 °C with 16 h/8 h light/dark cycles, LED illumination at 400 μmol.m^-2^.s^-1^, and 70% humidity. Plants were watered to maintain 70% soil moisture three times a week with osmosed water. Weekly, tomatoes were fertilized with horticultural fertilizer (50 mL at 2 g.L^-1^) (9:16:36 of N:P:K, MultiTec™). Tomato plants were harvested at 28 days after sowing, at the vegetative stage. Roots were gently washed with a brush and untangled for length measurements. For metabolomic and transcriptomic analyses, tissues were rapidly snap-frozen in liquid nitrogen before being finely ground and stored at –80 °C. Leaf areas without petioles were measured using a planimeter (Li-3100 C Area Meter, Li-Cor, Lincoln, Nebraska, USA). Tissues used for carbon, nitrogen, nitrate and lignin content measurement were dried in an oven at 70 °C for 1 week and then finely ground using an orbital mill (Pulverisette 6, Fritsch, France) before being stored at room temperature in a dry-preserving package. Resulting powder was used for carbon, nitrogen and lignin measurements.

### 2.5. Total nitrogen and carbon analyses

Total C and N contents in dry tissues were determined using 3mg of dry matter, according to the Dumas method, with an elemental auto-analyzer (Unicube, Elementar, Langenselbold, Germany).

### 2.6. Cell wall characterization

Lignin and its components were quantified according to Lu *et al.,* 2021. Briefly, cell wall residues (CWR) were extracted using successive boiling in solvent baths. Lignin content was quantified in percentage of CWR according to the Cysteine-Assisted Sulfuric Acid (CASA) method. The lignin composition of CWR was measured using the thioacidolysis method as described by Rolando *et al.,* 1992, and the released fragments were analyzed using GC-MS at the VIB Metabolomics Core as described in (Hoengenaert *et al.,* 2022). The composition of the lignin was analyzed as described in (Hoengenaert *et al.,* 2022). Briefly, 20 mg of stems CWR were cellulase-digested and resuspended in DMSO. Scans of HSQC NMR spectra (Heteronuclear Single-Quantum Coherence) ^1^H–^13^C short-range correlation experiments were acquired for 6h, followed by a second 54h acquisition for improve sensitivity.

### 2.7. Metabolites extraction

Frozen powders from leaves, stems and roots (100 mg FW) were extracted twice using maceration (16 h in the dark at 22 °C with slow vertical agitation) followed by sonication (37 kHz, 10 min at room temperature), each time with 800 µL of methanol/deionized water (80/20). The macerates were then centrifuged (10 min, 16100 g, 22 °C). The supernatants (1.6 mL) were collected, dried (5-7 h under vacuum, Concentrator Plus, Eppendorf) and then resuspended in 100 µL of methanol/deionized water (80/20). The extracts were filtered (0.2 µm, Minisart® RC4 membrane, Sartorius, REF17821) and stored at –20 °C. Extracts were added with taxifolin (285 µM) as an internal quantification standard.

### 2.8. Metabolomics analysis

Chromatographic analyses coupled with High Resolution Mass Spectrometry were performed on a Thermo Scientific Vanquish UHPLC system equipped with a binary pump, an autosampler, a temperature-controlled column (40 °C) and a Diode Array Detector. Extracted samples were separated on a XB-C18 Kinetex (150 × 2.1 mm, 2.6 µm) (Phenomenex Inc., Torrance, CA, USA) using a gradient of mobile phase composed of 0.1% formic acid in water (A) and 0.1% formic acid in methanol (B) at a flow rate of 200 µL·min^−1^. The separation of the compounds was performed using a 30 min mobile phase gradient (A: B; v/v). The elution method is as follows: (90:10) at 0 min, (90:10) up to 2 min, (70:30) up to 10 min, (5:95) up to 20 min remaining 5 min, (90:10) up to 26 min. The molecules are detected according to their absorbance at different wavelengths between 190 and 800 nm.

HRMS^1^ detection was performed on an Orbitrap IDX^TM^ (ThermoFisher Scientific, Bremen, Germany) mass spectrometer in positive and negative electrospray ionization (ESI) modes. The capillary voltages were set at 3.5 kV and 2.5 kV for positive and negative modes, respectively. The source gases were set (in arbitrary unit min^−1^) to 40 (sheath gas), 8 (auxiliary gas) and 1 (sweep gas) and the vaporizer temperature and ion transfer tube was respectively 320 °C and 275 °C. Full scan MS^1^ spectra were acquired from 120 to 1200 m/z at a resolution of 60,000. MS^2^ analysis was performed on the pooled-samples vial using the data dependent acquisition (DDA) mode and fragments were analyzed by Orbitrap at a resolution of 15000. For this analysis, the AcquireX data acquisition workflow was applied. The raw data were uploaded on the Compound discoverer 3.3 software (ThermoFisher Scientific, Bremen, Germany) to run the untargeted metabolomics workflow (peak detection, chromatogram alignment and peak grouping in features). In addition, the full MS^1^–MS^2^ dataset was exported as a mascot generic format (mgf) file and analyzed through the Sirius 5 software (Dührkop *et al.,* 2019). Briefly, the procedure implies the automatic annotation of molecular formula using SIRIUS and ZODIAC (Dührkop *et al.,* 2019; Ludwig *et al.,* 2019). Then, molecular structure prediction was proposed through CSI: FingerID (Hoffmann *et al.,* 2021). Finally, feature was assigned to chemical families using CANOPUS (Djoumbou Feunang *et al.,* 2016; Dührkop *et al.,* 2019; H. W. Kim *et al.,* 2021). Further annotation was performed using Compound Discoverer 3.3 by predicting elemental composition, searching in local and public mass and formula databases such as ChemSpider, HMDB, and LipidMaps, and querying spectral databases including mzCloud, MoNA, and GNPS.

Formal identification of coumarins and furanocoumarins, was afforded via comparisons with commercial standards according at least to their retention time (RT), mass (MS^1^) and mass fragmentation (MS^2^) when available. A standard curve was established for the scopoletin quantification using as internal standard the taxifolin (285 µM).

### 2.9. Construction and analysis of RNAseq libraries

Total RNA from leaves and roots was extracted from 3 plants per genotype (WT and PP) as previously described (Roumani *et al.,* 2022). RNA concentrations were determined from their absorbance at 260 nm. Their purity was evaluated by the 260/280 nm absorbance ratio, their quality was checked by electrophoresis on a 1% agarose gel and their integrity by determining and considering an RNA Integrity Number (RIN) >7 (Agilent BioAnalyzer 2100). Transcript data was obtained by RNA-Seq Illumina pair-end sequencing (2× 150pb) using the NovaSeq 6000 methods for 30 million read pairs. Raw data were cleaned using Trimmomatic (Bolger *et al.,* 2014). The quality of the sequences was assessed using FastQC. The transcripts were mapped on the tomato genome (ITAG2.4) using the BWA-MEM software (Li & Durbin, 2009). Count tables were generated using the FeatureCounts software (Version 1.6.4) and Samtools merge workflows (Liao *et al.,* 2014). The differential gene expression was performed with the glmfit option within the edgeR package for a FDR <0.05 (Robinson *et al.,* 2010) and using AskoR (Alves-Carvalho *et al.,* 2021). We performed, within AskoR, a clustering using kmeans classification algorithms (Coseq R package) (Godichon-Baggioni *et al.,* 2019). Functional annotations, such as gene ontologies and interproscan features, were extracted from ITAG2.4_protein_functional.ge3 file. Differentially expressed genes (DEGs) between WT and PP lines DEGs were used to perform GO enrichment analysis (GOEA) restricted to the Biological Processes on ShinyGO (version 0.80) using the “*Solanum lycopersicum* genes SL3.0” and a FDR cutoff <0.05 (Ge *et al.,* 2020). RNA-seq dataset from tomato leaves and roots is publicly available through the Data.gouv.fr repository: https://doi.org/10.57745/G2Z4VV.

The results of the RNAseq differential analysis were confirmed by following the expression level of 2 genes (*Sl*COSY and *Sl*PRCA) in addition to the 5 engineered genes in leaves and roots of the PP and control tomato plants. The list of targeted genes and primers used is given in **Table S2h**. The expression level was followed by RT-qPCR. For that purpose, a total of 200 ng of DNA-free RNA samples was reverse transcribed in a final volume of 20 µL using the RNA to cDNA kit (Life technologies, Tokyo, Japan). cDNAs were then diluted 10 times. Gene expression was measured by qPCR on a CFX Duet Realtime PCR System (Bio-Rad). Each PCR reaction was made in 20 µL containing primer pairs at 0.3 µM, 40 ng of cDNAs in 1× Mesa Blue Premix (Eurogentec) and followed the amplification protocol consisting in a denaturing step at 95 °C for 30 s, 40 cycles at 95 °C for 5 s and 60 °C for 30 s. The gene expression of each sample was calculated with three analytical replicates and normalized to the two internal control genes *ACT* and *GAPDH*.

### 2.10. Statistical Analysis

Measured and calculated morphometric and physiological variables, as well as relative and absolute quantified metabolites were statistically analyzed by comparing WT and PP lines using a non-parametric Kruskal-Wallis test.

## 3. Results

### 3.1. The PP line regeneration

To insert the Psoralen Pathway (PP) in tomato plants, we first leveraged the GoldenBraid technology to assemble the selected four genes encoding the enzymes required for the conversion of *p*-coumaroyl-CoA into psoralen. The T-DNA was build-up with 5 genes organized as transcription units (TUs) (Larbat *et al.,* 2009; Munakata *et al.,* 2016; Roselli *et al.,* 2017; Villard *et al.,* 2021), each independently under the control of the constitutive *CaMV*35S promoter and the tNOS terminator (**Fig. 1**; **Fig. S1**). Then, we integrate the PP T-DNA into the genome of the *Wva*106 tomato cultivar using an *agrobacterium*-mediated stable transfection. Plant selection was allowed by a fifth similarly assembled TU including the kanamycin resistance. By assessing genomic integration via PCR on gDNA, we successfully regenerated two independently transfected lines (**Fig. S1b**). Among the two lines, qPCR revealed that only one line (the PP line) showed a constitutively high expression levels of each transgene in the investigated leaves and roots, compared to the WT (**Table S2g,h; Fig. S1c**). Our study focused on this particular line, which exhibited a phenotype similar to the WT (**Fig. S1e**).

### 3.2. The PP line accumulates coumarins

To investigate the impact of Psoralen Pathway integration on tomato plant metabolic composition, we compared the metabolomes of PP and WT plants in leaves, stems, and roots using untargeted LC-HRMS analysis of hydro-methanolic extracts. To strengthen the analysis, given that only one PP line was obtained, we performed three independent cultivation rounds under identical growing conditions, resulting in 23 WT and 19 PP plants. The metabolome comparison between PP and WT in all the three-round cultivation of plants and across the three organs, highlighted an overall tendency to a higher accumulation of dozens to hundred metabolic features in the PP line. By considering only significant metabolic features that differentially accumulated (Benjamini-Hochberg p<0.05), 82 metabolic features across leaves, stems and roots were retained (**Table S1**). One metabolic feature was less abundant in the PP line whereas all others were enriched in comparison to the WT. Putative annotation of these features allowed to categorize 20 of them as fatty acids and 19 as phenylpropanoids (*i.e.,* 10 coumarins, 7 phenolamides, the furanocoumarin: psoralen, and one miscellaneous compound) (**Fig. 2a**). The abundance of only eight of the metabolic features were significantly affected in at least two organs, and among them scopoletin, the only metabolite that was found to be differential in leaves, stems and roots (**Fig. 2b**).

**Figure 2:**
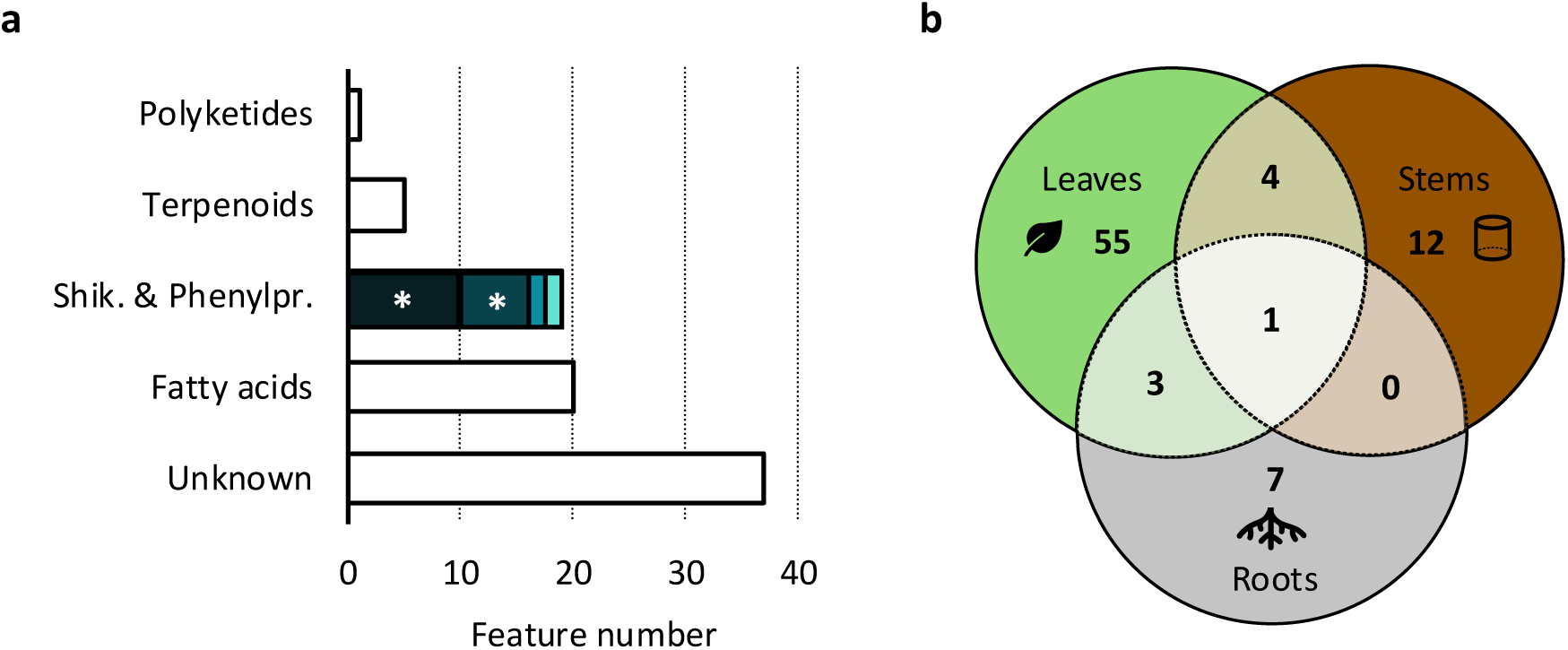
Untargeted comparative metabolomic profiling of the Psoralen Pathway (PP) and the Wild-Type (WT) tomato lines. (**a**) Number of metabolic features significantly over-accumulated in the PP line compared to the WT. Metabolic features are categorized according to the NP-Classifier ontology (Kim *et al.,* 2021). Shikimate and phenylpropanoids are highlighted in blue from dark to light: coumarins*, phenolamides*, others, furanocoumarins. Asterisks* correspond to metabolic subclasses significantly enriched (**Table S1b**). (**b**) Venn diagram describing the distribution of the PP line’s over-accumulated features across leaves (green), stems (brown) and roots (grey) compared to the control line. Number of replicates for leaves, stems, roots: WT (n=23), PP (n=19). The data are based on 3 independent cultivation rounds of tomato plants grown in the same phytotronic conditions.

We then performed targeted analyses focusing on coumarins and metabolites of the psoralen pathway. Psoralen and its precursors (*i.e.,* demethylsuberosin and marmesin) were detected only in trace amounts across the analyzed organs of the PP line, and were absent from the WT (**Fig. 3a**). The PP line exhibited significant increases in *p*-coumaric acid and umbelliferone levels in the leaves (50– and 31-fold, respectively in comparison to WT), with no significant changes in other plant parts. Scopoletin concentrations were significantly increased in all organs of the PP line (23-to 1799-fold), and especially in stems where it reached concentrations close to 5 µg/g of DW (**Fig. 3b**). Esculetin and its glycoside form esculin accumulated significantly higher in the PP line than in the WT (**Fig. 3a**). On the opposite, scopolin accumulation was not different between the WT and the PP line.

**Figure 3:**
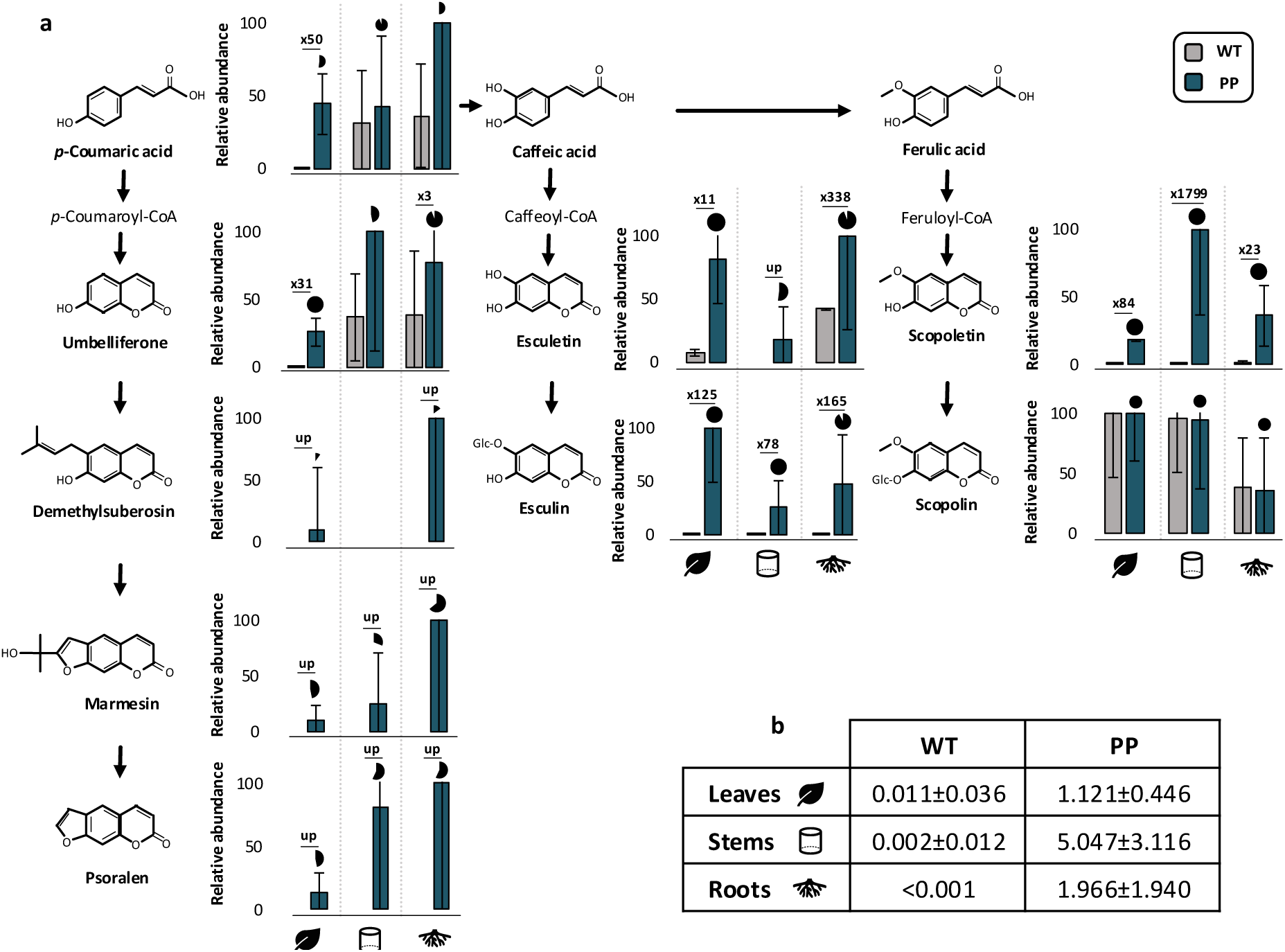
Comparative targeted profiling of coumarins and furanocoumarins in the Psoralen Pathway (PP) and the Wild-Type (WT) tomato lines. (**a**) Relative accumulation of reported coumarins and furanocoumarins in the PP line (blue) versus the WT line (grey), expressed as mean ± standard deviation. Percentage of replicates containing the metabolite in the PP line is illustrated using black pie charts. Only significantly dicerentially accumulated metabolites, assessed by the Kruskal-Wallis test (p<0.05*), are depicted according to their increasing fold change in the PP line compared to the WT line. ‘Up’ refers to compounds detected only in the PP line. (**b**) Absolute quantification of scopoletin in µg/g of dry weight. Data are expressed as percentages (a) or absolute quantity for each metabolite relative to the most abundant peak area, normalized to the internal standard taxifolin, and reported per mg of dry weight; Number of replicates for leaves, stems, roots: WT (n=23), PP (n=19). The data are based on 3 independent cultivation rounds of tomato plants grown in the same phytotronic conditions.

### 3.3. Scopoletin is released by a lignin-degradative method

Scopoletin has recently been described to be able to incorporate into the lignin polymer of *Arabidopsis thaliana* (Brassicaceae) and *Populus sp* (Salicaceae) transgenic lines producing this metabolite in the stems (Hoengenaert *et al.,* 2022; N. Wang *et al.,* 2025). Accordingly, the lignin composition was analyzed in the PP line and compared to that in the WT. Lignin content (**Fig. 4a**) and composition (H, G, S) (**Fig. 4b-c**) in the stems of the PP line were not significantly affected compared to WT. However, analysis of the thioacidolysis released monomers indicated a slight but significant increase in the release of scopoletin in the PP line (2.8×) compared to the WT line (**Fig. 4c**), suggesting the presence of scopoletin in the cell wall as well as in the lignin.

**Figure 4:**
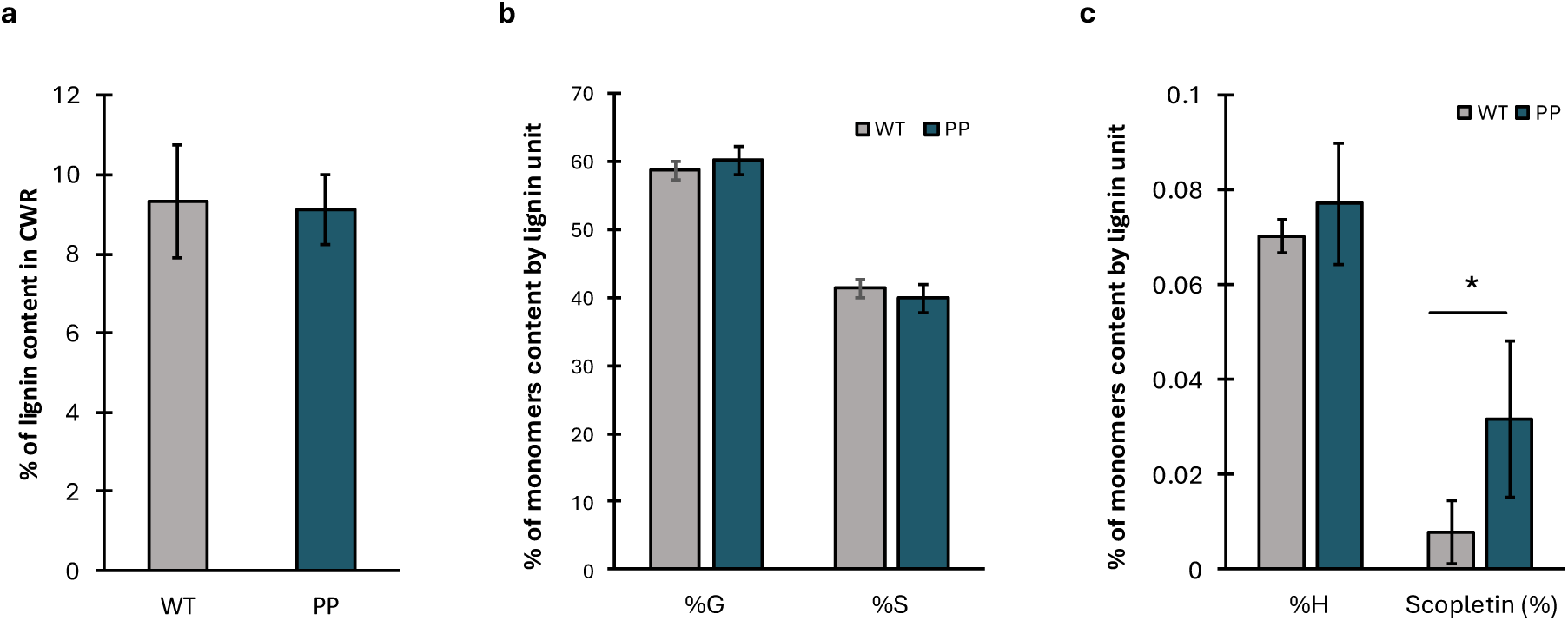
Impact of the Psoralen Pathway (PP) line on lignin content (a) and composition for G, S (b), and H monomers, as well as for scopoletin (c), in stems compared to the Wild-Type (WT) line. The average percentage of monomers and scopoletin are calculated based on the content of H, G, and S units and scopoletin in the lignin. The error bars indicate the standard deviation. The lignin content was measured as percentage of Cell Wall Residue (CWR) using the Cysteine-Assisted Sulfuric Acid (CASA) method (Lu *et al.,* 2021). Data were compared using the Kruskal-Wallis’s test (p-value<0.05) (p-value for scopoletin % = 0.00106); WT (n=8) and PP (n=9).

### 3.4. Specific genes regulation in the PP line

To further understand the metabolic changes observed in the PP line, we then investigated its transcriptome compared to the WT. Results highlighted 1014 and 53 differentially expressed genes (DEGs) in the leaves and roots, respectively, of the PP line (**Fig. 5a; Table S2**). In leaves, the 1014 DEGs comprised 659 upregulated and 355 downregulated genes in the PP line compared to WT. In roots, 41 genes were upregulated and 12 downregulated. Seventeen DEGs were common to leaves and roots. Gene Ontology Enrichment Analysis (GOEA) was performed on the up– and downregulated DEGs identified. In leaves, GOEA analysis highlighted an overall upregulation of genes involved in processes related to response to stress and wounding (**Fig. 5b**), whereas the main trend was a downregulation of genes associated to chlorophyll biosynthesis (**Fig. 5c**). In roots, the relatively short list of DEGs did not lead to any significant enrichment using GOEA.

**Figure 5:**
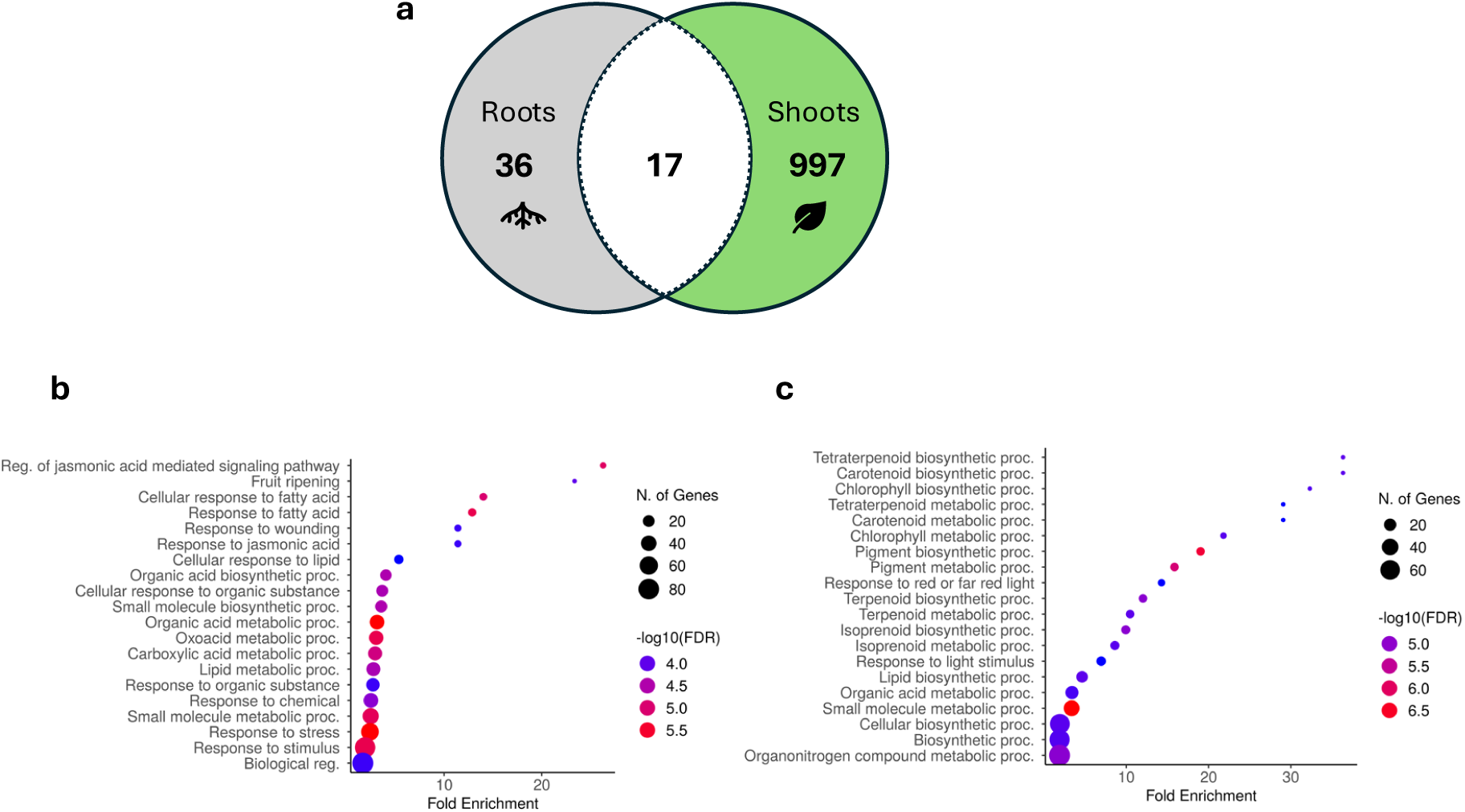
Transcriptomic regulation in the Psoralen Pathway (PP) line compared to the Wild-Type (WT) line. (**a**) Dicerentially Expressed Genes (DEGs) in leaves and/or roots (FDR<0.05). Gene Ontology Enrichment Analysis (GOEA) on biological processes for (**b**) upregulated and (**c**) downregulated genes considering only leaves (False Discovery Rate: FDR<0.05). The position of the circles represents the fold change by related to the ratio PP/WT. The size of the circles represents the number of Dicerentially Expressed Genes (DEGs) in each Gene Ontology (GO) category. The color of the circles indicates the statistical significance. The data consist of 3 replicates of PP and WT per tissue.

Using a differential gene analysis based on 3 replicates in leaves and roots comparing the PP and WT lines, we highlighted only strongly DEGs in the PP line (**Fig. 5**). Finer regulatory patterns were uncovered through a K-means clustering approach, which revealed 11 expression clusters based on gene expression profiles in leaves and roots, comparing the PP and WT lines (**Fig. S2**). Among them, five clusters displayed expression patterns that distinguished the PP line from the WT. The smaller and most specific expression cluster was cluster 8 regrouping 309 genes with a higher expression in both the leaves and roots of the PP line in comparison to WT. In addition, cluster 2 (1449 genes) contained genes specifically over-expressed in the roots of the PP line, and cluster 7 (1113 genes), over-expressed genes in the leaves. Conversely, the clusters 1 (743 genes) and 5 (1466 genes) grouped genes downregulated in leaves and roots of the PP line compared to the WT. GOEA of cluster 8 indicates the induced expression of genes potentially linked to the PP line’s response to stress, likely signaled *via* jasmonate and in response to lipids (**Fig. 6a**). As an extension of cluster 8, the same trend was observed as for the genes from cluster 7 (upregulated in leaves of the PP line) (**Fig. 6b**). The cluster 2 (upregulated genes in roots of the PP line), were enriched in processes related to hormonal regulations (**Fig. 6c**). Among the processes related to downregulated genes (**Fig. 6d,e**), DNA replication and cell cycle appeared affected in the roots of the PP line (cluster 5), whereas lipid and sterol biosynthesis were, in leaves (cluster 1).

**Figure 6:**
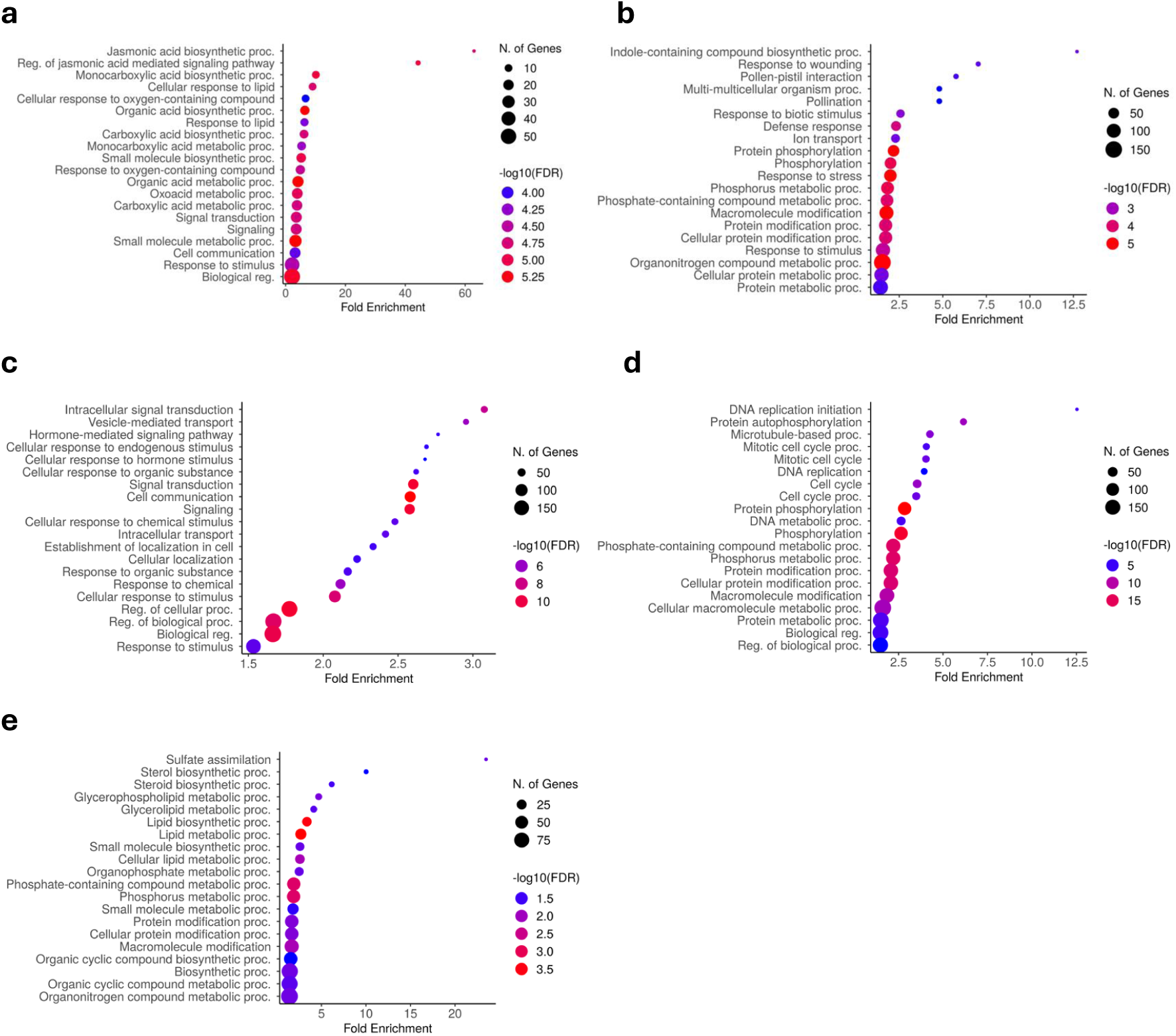
Gene Ontology Enrichment Analysis (GOEA) on biological processes in the Psoralen Pathway (PP) line compared to Wild-Type (WT) line for clusters 8 (a), 7 (b), 2 (c), 5 (d) and 1 (e). (*False Discovery Rate:* FDR<0.05). The position of the circles represents the fold change by related to the ratio PP/WT. The size of the circles represents the number of Dicerentially Expressed Genes (DEGs) in each GO category. The color of the circles indicates the statistical significance. Gene clusters were grouped using the k-means method. (**a**) Cluster 8 includes 309 upregulated genes and most of those were specifically dicerentially expressed in the leaves and roots of the PP line compared to the WT line (10/17 genes). (**b**) Cluster 7 contains 1113 upregulated genes in leaves. (**c**) Cluster 2 contains 1449 upregulated genes in roots. (**d**) Cluster 1 and (**e**) cluster 5 contain 743 and 1465 downregulated genes in roots and leaves, respectively. The data consist of 3 replicates of PP and WT per tissue.

### 3.5. The PP line showed reduced biomass and elevated nitrogen content

To identify potential impacts on the growth and development of the PP line compared to the WT line, we performed a comparative resource allocation and morphology analysis (**Table S3**). After 28 days of growth, PP and WT plants exhibited comparable morphology and size (**Fig. S1e**), with similar leaf numbers and stem lengths between days 13 and 28, indicating no significant developmental changes (**Table 1**). However, dry biomass in the PP line was significantly reduced by 21.3% compared to the WT. This decrease was uniformly distributed across plant organs with the exception of stem which had a similar amount of dry biomass compared to the WT. As a results, the resource allocation pattern in the PP line was significantly different from the WT, with a decrease in the proportion of matter in the leaves (−10%) and an increase in the stem (+20%) (**Table 1**). The decrease in the biomass accumulation in leaves was in accordance with an increase in the Specific Leaf Area (SLA) in the PP line.

**Table 1:** Mean alterations in growth and development parameters between WT and the PP line. The significant dicerences between the two genotypes are supported by a non-parametric Kruskal-Wallis test. Significant dicerences are indicated in bold with an asterisk. Measurements were taken at a vegetative growth stage (28 days after sowing; D28) and compared for development with measurements at an early growth stage (13 days after sowing; D13). Dry biomass allocation patterns are defined by the percentage of mean dry biomass in each compartment. Abbreviations: SLA (Specific Leaf Area); SRL (Specific Root Length); SSL (Specific Shoot Length). WT (n=23), PP (n=19).

| | Variable | WT average ± SE | PP average ± SE | p-value |
| --- | --- | --- | --- | --- |
| Dry weights | Foliar blades (g) | 2.46 ± 0.36 | 1.74 ± 0.24 | **0.0004\*** |
| | Petioles (g) | 0.49 ± 0.07 | 0.4 ± 0.05 | **0.0222\*** |
| | Stems (g) | 1.11 ± 0.28 | 1.05 ± 0.17 | 0.4142 |
| | Roots (g) | 0.49 ± 0.05 | 0.39 ± 0.06 | **0.0025\*** |
| | Stems/Roots ratio | 8.20 ± 1.07 | 8.15 ± 0.70 | 0.9349 |
| Tissue allocation<br>(% by DW) | Foliar blades | 54.06 | 48.6 | **0.0042\*** |
|  | Petioles | 10.76 | 11.17 | 0.2529 |
|  | Stems | 24.39 | 29.32 | **0.0178\*** |
|  | Roots | 10.76 | 10.89 | 0.9349 |
| Development | Stems (cm) | 49.95 ± 7.42 | 51.33 ± 6.02 | 0.6532 |
| | Roots (cm) | 44.21 ± 15.1 | 44.22 ± 9.38 | 0.7439 |
|  | Leaf number | 8 | 8 | 1 |
| | D13/D28 plant height | 0.1 ± 0.02 | 0.1 ± 0.02 | 0.8064 |
| Key growth analysis parameters | Stems/Roots ratio | 1.24 ± 0.43 | 1.2 ± 0.28 | 1 |
|  | SLA (cm<sup>2</sup>.g<sup>-1</sup>) | 236.48 ± 33.67 | 329.73 ± 35.81 | **0.0002\*** |
|  | SRL (cm.g<sup>-1</sup>) | 90.95 ± 39.82 | 115.21 ± 31.85 | 0.0724 |
|  | SSL (cm.g<sup>-1</sup>) | 45.06 ± 14.62 | 47.84 ± 6.17 | **0.0274\*** |

To better understand the cause of the loss in biomass in the PP line, we analyzed carbon (%C) and nitrogen (%N) accumulation across the different tissues (**Table 2**). Results showed that nitrogen content was significantly higher in the PP line compared to the WT, particularly in the shoots (+53%). Focusing on ion uptakes in leaves and roots (**Table S4**), we identified a significant increase of nitrate ions in roots (3.3×). Conversely, carbon content was slightly lower in the PP line, most notably in the stems (−13%). As a result, the carbon-to-nitrogen ratio (C/N) was significantly decreased in the PP line, especially in the shoots.

**Table 2:**
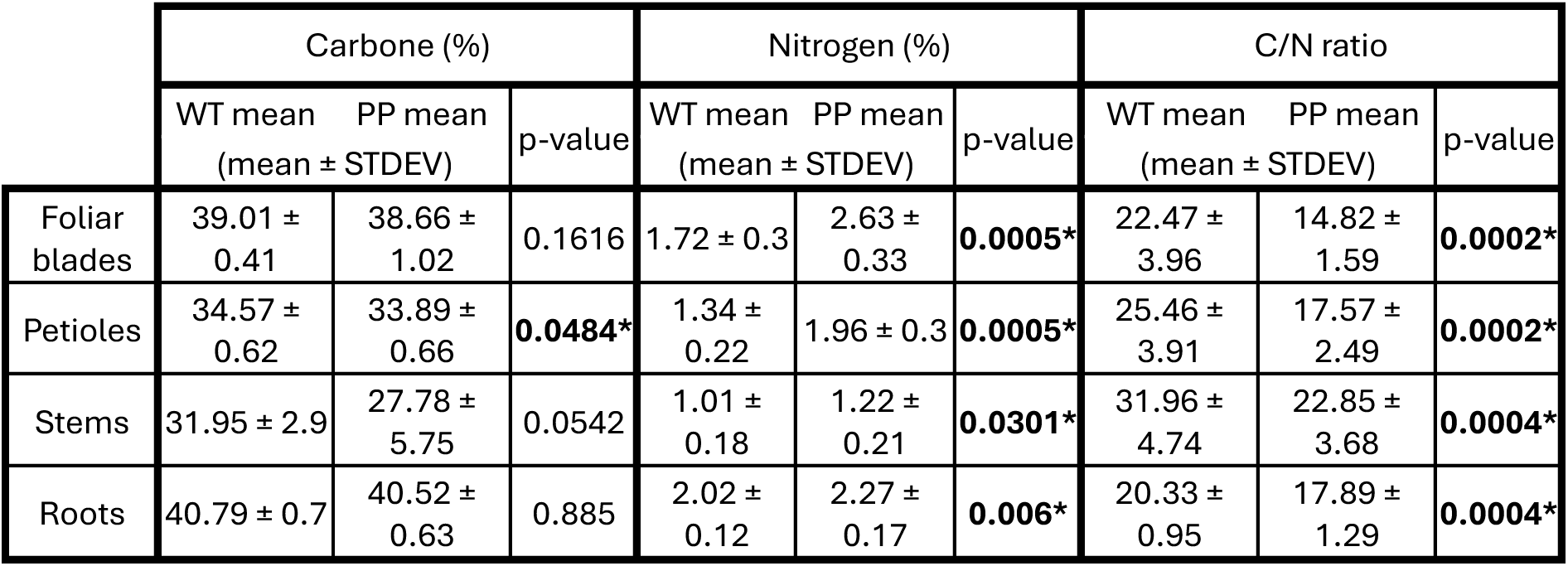
Mean alterations in Carbon (C), Nitrogen (N) contents and C/N ratio between the Wild-Type (WT) and the Psoralen Pathway (PP) lines. The variables significance between the two genotypes is supported by a non-parametric Kruskal-Wallis test. Significant dicerences are indicated in bold with an asterisk. Measurements were taken at a vegetative growth stage (28 days after sowing). WT (n=10), PP (n=9).

|  | Carbone (%) |  |  | Nitrogen (%) |  |  | C/N ratio |  |  |
| --- | --- | --- | --- | --- | --- | --- | --- | --- | --- |
|  | WT mean<br>(mean ± STDEV) | PP mean<br>(mean ± STDEV) | p-value | WT mean<br>(mean ± STDEV) | PP mean<br>(mean ± STDEV) | p-value | WT mean<br>(mean ± STDEV) | PP mean<br>(mean ± STDEV) | p-value |
| Foliar blades | 39.01 ± 0.41 | 38.66 ± 1.02 | 0.1616 | 1.72 ± 0.3 | 2.63 ± 0.33 | **0.0005\*** | 22.47 ± 3.96 | 14.82 ± 1.59 | **0.0002\*** |
| Petioles | 34.57 ± 0.62 | 33.89 ± 0.66 | **0.0484\*** | 1.34 ± 0.22 | 1.96 ± 0.3 | **0.0005\*** | 25.46 ± 3.91 | 17.57 ± 2.49 | **0.0002\*** |
| Stems | 31.95 ± 2.9 | 27.78 ± 5.75 | 0.0542 | 1.01 ± 0.18 | 1.22 ± 0.21 | **0.0301\*** | 31.96 ± 4.74 | 22.85 ± 3.68 | **0.0004\*** |
| Roots | 40.79 ± 0.7 | 40.52 ± 0.63 | 0.885 | 2.02 ± 0.12 | 2.27 ± 0.17 | **0.006\*** | 20.33 ± 0.95 | 17.89 ± 1.29 | **0.0004\*** |

## 4. Discussion

The biosynthesis of specialized metabolites necessarily triggered a physiological cost on plants, either by diverting resources for their production or through the pleiotropic effects of the specialized metabolites themselves (Neilson *et al.,* 2013). This study aimed to better understand the impacts of constitutive expression of four genes from the linear furanocoumarin biosynthesis pathway on tomato plant physiology and growth.

### 4.1. Furanocoumarins might be toxic for tomato plants

Genetic modification of tomato with our construct aimed at producing psoralen surprisingly resulted in a low transformation success rate (0.33%), especially when compared to longer pathways already inserted with success in tomato using the same GoldenBraid system with TUs also under the *CaMV*35S promoter (H. Kim *et al.,* 2025; Sonawane *et al.,* 2017). Furthermore, psoralen was found only as trace amounts despite that all PP transgenes were constitutively expressed (**Fig. S1c**; **Table S2g**). In parallel, transcriptomic data showed significantly over-expressed genes falling in the GO categories “response to stress”, “response to chemicals” and other direct or indirect stress-related GO classifications in the PP line (*e.g.,* photoinhibition, stress phytohormones, etc) (**Fig. 5b,c**; **Fig. 6a-c**). These findings suggest a psoralen pathway-dependent phytotoxicity in tomato plants, potentially linked to the production of furanocoumarins (**Fig. 3a**). During the seven last decades, incremental progress has highlighted a large spectrum of furanocoumarin’s toxicity on a vast range of organisms, including plants, occurring through a diversity of processes (*e.g*., photocycloaddition, lipid peroxidation, cytochrome P450 inhibition) (Baskin *et al.,* 1967; Bennet & Bonner, 1953; Diawara *et al.,* 1993; Frank *et al.,* 1998; Greenberg *et al.,* 2001; Hale *et al.,* 2004; Hassan *et al.,* 2016; Kitamura *et al.,* 2005; Meepagala *et al.,* 2021; Razavi, 2010; X.-B. Wang *et al.,* 2008). We therefore hypothesize that psoralen, or core intermediates and/or derivatives of its biosynthetic pathway, may be toxic, or even lethal, to tomato above a certain threshold. This toxicity may have impaired the regeneration of additional transgenic tomato lines. However, although our transcriptomic data clearly indicated a chemically related phytotoxicity in the PP line, no clear signal allowed us to pinpoint if this toxicity was related to the production of psoralen and/or any intermediates, or even to the accumulation of any of the other accumulated compounds, in particular coumarins. However, the fact that the PP line plants appeared healthy (**Fig. S1e**) suggests that they were able to circumvent the induced phytotoxicity.

### 4.2. The low accumulation of furanocoumarins may reflect metabolic competition favouring the coumarin biosynthesis pathway

Metabolic engineering approaches often fall short of expectations due to our limited understanding of the self-management of specialized metabolite biosynthetic pathways in plants (Lynch *et al.,* 2021). In the PP line, psoralen as well as its precursors from umbelliferone could be detected in almost all the plant parts but at a level below the quantification limit (**Fig. 3a**), proving that our metabolic engineering approach is working but not sufficiently well to allow psoralen accumulation. Given that all genes encoding the enzymes essential for psoralen accumulation were appropriately expressed and the corresponding enzymes proven functional (**Fig. S1e**; **Table S2g**), we suggest a metabolic competition for its precursor, *p*-coumaroyl-CoA, which may be steered instead toward coumarins, which were the more enriched in the PP line. Indeed, simple coumarins like umbelliferone, esculetin, and scopoletin derive from *p*-coumaroyl-, caffeoyl-, and feruloyl-CoA, respectively, via 2OGD-mediated *ortho*-hydroxylation followed by a cyclisation that occurs spontaneously and/or is catalyzed by a COSY (C. Y. Kim *et al.,* 2023; Vanholme *et al.,* 2019) (**Fig. 1**). The *ortho*-hydroxylation of *p*-coumaroyl-CoA constitutes the first intended metabolic step for psoralen production in the PP line. To allow umbelliferone accumulation for psoralen production, we chosen the C2′H 2OGD from *Pastinaca sativa* (Apiaceae) (*Ps*Diox) which was characterized *in vitro* as being specific for *p*-coumaroyl-CoA (Roselli *et al.,* 2017). The marked and significant accumulation of coumarins in the PP line including the formally identified esculin and scopoletin (**Fig. 3**, **Fig. S3**), suggests that, *in vivo*, *Ps*Diox could share the same catalytic properties as its orthologs from *Ruta graveolens* (Rutaceae), *A. thaliana* and *Ipomea batatas* (Convolvulaceae) that also accept caffeoyl-CoA and feruloyl-CoA as substrates (Matsumoto *et al.,* 2012; Vialart *et al.,* 2012; Yang *et al.,* 2015). This hypothesis is supported by the accumulation of scopoletin and esculetin in plants overexpressing F6′H (Beesley *et al.,* 2023; N. Wang *et al.,* 2025).

Non-exclusive hypotheses for the limited accumulation of psoralen may also involve active regulation of competing metabolic pathways that bypass the psoralen branch (**Fig. 3**). Indeed, comparative metabolomics analysis highlighted the accumulation of phenolamides (*i.e.,* fused hydroxycinnamates and amines), a natural defence compound family in tomato plants (Roumani et al., 2022), in addition to coumarins (**Fig. 2**, **Table S1**). In parallel, comparative transcriptomics pointed to the overexpression in the PP line of genes of the core phenylpropanoid pathway putatively involved in the coumarin and phenolamide biosynthesis pathways (*e.g*., BAHD acyltransferases, Tyramine N-feruloyl transferase) (**Table S2c**). Such metabolic competition for a common substrate such as *p*-coumaroyl-CoA could also be further established by other factors like metabolic channeling as now established in several species (Dahmani *et al.,* 2023; Lallemand *et al.,* 2013; Laursen *et al.,* 2016). Heterologous enzymes from different plant species may be less competitive than endogenous ones in establishing metabolic channeling of intermediates and co-substrates, ultimately at the expense of psoralen accumulation.

### 4.3. Coumarin accumulation might underly the PP line’s altered physiology and growth

The metabolomics analyses highlighted 82 metabolic features almost exclusively over-accumulated in the transgenic line. Annotation outlined that primarily 4 metabolic families (fatty acids, phenolamides, coumarins and furanocoumarins) were involved. Among them, coumarin accumulation stood out from the other families for at least one of the following reasons: (i) the accumulation of all individual coumarins followed a presence/absence pattern in the PP/WT comparison (**Fig. 3a**; **Table S1**), (ii) the accumulation of coumarin metabolites was detected in the three tissues (leaf, stem, roots) in accordance to the constitutive expression of the PP transgenes (**Fig. 3a**) and (iii) based on the targeted analyses the accumulation of coumarins reached several µg/g of DW (*i.e.,* scopoletin) whereas the furanocoumarin accumulation did not reach the quantification limit (**Fig. 3**). These observations made us consider that coumarins accumulation may be the main stimulus of the transcriptomic, physiological and developmental characteristics of the PP line. This would be consistent with previous studies in which coumarins (*i.e.,* umbelliferone, scopoletin, coumarin, and daphnoretin) were applied to *A. thaliana* and *Lactuca sativa* (Asteraceae), resulting in allelopathic effects such as membrane peroxidation, Reactive Oxygen Species (ROS) production, mitotic disruption, and photoinhibition (Araniti *et al.,* 2017; Graña *et al.,* 2017; Z. Yan *et al.,* 2016). Our transcriptomics analysis highlighted the upregulation, in the PP line in comparison to WT, of a majority of genes associated with responses to stress, chemicals, organic substances and wounding (**Fig. 5b**; **Fig. 6a-c**). These categories comprise genes involved in ROS detoxification (*e.g.,* glutathione-S-transferases, cytochrome P450s, peroxidases, detoxification proteins, chaperones, and heat-shock proteins) (**Table S2**). On the contrary, the main categories of downregulated genes in the leaves of the PP line compared to WT were associated to chlorophyll biosynthesis (**Fig. 5c**; **Fig. 6d**). These transcriptomics patterns suggest an active regulation of the plant oxidative status, potentially mediated by an increase in the ROS detoxication processes and a decrease in the ROS production linked to photosynthesis. This suggestion is supported by the healthy appearance of the PP line despite the activation of all these stress signaling pathways (**Fig. S1e**). As now well established, photoinhibition generally acts as a protective mechanism to limit photoinduced ROS accumulation, as reviewed by Didaran *et al.,* 2024. In parallel, the lipid over-accumulation observed in the leaves of the PP line (**Table 1**), together with the over-expression of genes associated with lipid metabolism (*e.g.,* phospholipases, aldehyde dehydrogenases, lipid biosynthesis) and membrane repair (*e.g.,* lipid desaturases, acyltransferases) (**Fig. 6a**; **Table S2**), further supports the occurrence of membrane damage. This lipid stress may result from coumarin-induced destabilization of chloroplast integrity and photochemistry, as previously observed in *A. thaliana* (Araniti *et al.,* 2017).

To further understand the impact of coumarins on tomato physiology in the PP line, we investigated its effects on growth, development and resource allocation compared to the WT line. First, we observed no developmental alterations in the PP line, in contrast to the pronounced growth impairments reported in various plant species metabolically engineered to over-accumulate scopoletin (Beesley *et al.,* 2023; Hoengenaert *et al.,* 2022; N. Wang *et al.,* 2025). However, F6′H knock-down lines of *Manihot esculenta* (Euphorbiaceae) with reduced scopoletin accumulation have revealed key roles for this compound in cassava development (*e.g*., auxiliary budding, leaf appearance, apical dominance) (Mukami *et al.,* 2024). All these elements suggests that the physiological effects of coumarins on development may depend on their constitutive accumulation levels and may be species-specific. This also underscores how poorly understood the physiological regulation of coumarins remains, despite their strong potential to act in a phytohormone-like manner, as hypothesized for an auxin-like effect of scopoletin by Graña *et al.,* 2017.

Morphological analysis of the PP line revealed a significant biomass reduction, particularly of the leaves (**Table 1**). A similar biomass loss was previously reported in *A. thaliana* supplemented with coumarin or scopoletin (Araniti *et al.,* 2017; Graña *et al.,* 2017) and was partly attributed to photoinhibition observed in the leaves, similarly to what we found in the PP line according to transcriptomic data (**Fig. 6d**). In addition, we also observed a downregulation of a few genes putatively involved in sugar storage (*i.e.,* starch synthase) and numerous genes putatively involved in sugar metabolism and cell wall remobilization (**Table S2c,d**). Their differential expression may help explain the biomass loss observed, which cannot be exclusively attributed to carbon content reduction, as this decrease is substantially lower than the 21% drop in biomass (**Tables 2 and 3**). Considering that most of the biomass components were measured (*e.g.,* carbon, nitrogen, ions) (**Table 2**; **Table S4**), we suggest a concomitant reduction in oxygen-rich compounds such as sugars. In parallel, significant increase of specific leave surface (SLA; +39%) underlies a biomass loss that is distributed within thinner leaves (**Table 1**). Taken together, these elements suggest that coumarins may trigger a ‘slimming effect’, leading to biomass reduction, primarily in tomato leaves, as a potential response to coumarin-induced phytotoxicity.

Thioacidolysis revealed that scopoletin was incorporated into the cell wall (**Fig. 4c**), albeit at a level approximately 10 to 100 times lower than found in *A. thaliana* and *Glycine max* (Fabaceae) (Beesley *et al.,* 2023; Hoengenaert *et al.,* 2022). Although only trace amounts of scopoletin accumulated in the cell wall, which prevented validation of its integration into the lignin polymer by NMR, it is reasonable to consider that a small fraction of scopoletin could be incorporated into the lignin polymer. Although the stem cell wall appears to be a key storage compartment for coumarins (Beesley *et al.,* 2023), glycosylated coumarins are stored in vacuoles, *e.g.,* in *A. thaliana* roots (Robe *et al.,* 2021). Both vacuolar storage and cell wall transfer are well established toxicity mitigation processes (Weng *et al.,* 2021). Our metabolic and phenotypic data suggest that even a modest accumulation of scopoletin (∼1-5 µg/g DW), has to be mitigated in the PP line.

Finally, we found that the significant increase in nitrogen content in the upper parts of the PP line coincided with enhanced nitrate uptake by the roots. Catecholic coumarins such as esculetin and fraxetin are well-known to facilitate iron uptake from the soil in plants (H. H. Tsai & Schmidt, 2017). Our data suggest that coumarins accumulation, primarily esculetin, is concomitant with nitrate uptake, highlighting a potential role of coumarins in this process. A similar contribution of coumarins already reported in *Triticum durum* and *Zea mays* (both Poaceae) (Abenavoli *et al.,* 2001; Lupini *et al.,* 2018).

## 5. Conclusion

The production of psoralen in tomato proved to be more challenging than the mere expression of the biosynthetic genes as it potentially led to pronounced phytotoxicity. Our results suggests that the accumulation of coumarins at concentrations of 1-5 µg/g of DW in tomato plants could induces several levels of regulation: (i) xenobiotic stress leading to oxidative stress, which the plant can adapt; (ii) a photoinhibition and an increase in metabolism inducing a biomass slimming possibly in response to oxidative stress; and (iii) a modification of the nitrogen and nitrate levels. Our results indicate that, although coumarins hold substantial agronomic potential (non-toxic to pollinators, facilitating iron uptake, shaping root microbiota, etc) (Cosme et al., 2021; H.-H. Tsai et al., 2018; Zhou et al., 2023), their role in plant defense entails a growth cost even at low content. A counterpart that should be carefully considered and ideally mitigated, following a more precise delineation of the regulatory pathways of coumarins underlying the identified physiological impacts, before pursuing potential viable applications.

## 6. Acknowledgments

We would like to thank Alexandre Olry, Claude Gallois and David Marcolet members of the Experimental phytotronic platform of Lorraine PEPLor (Université de Lorraine, France) for their technical assistance, Cédric Paris from the Metabolomic and Structural Analytic Platform facility (PASM, Université de Lorraine, France) as well as Geert Goeminne and Sandrien Desmet from the Metabolomic Core (VIB, Gent, Belgium). We thank Sarah Liu for her assistance on NMR sample preparation. We thank Florent Magot (BBV, Université de Tours) and Florent Ducrocq for assistance in the critical analysis of the metabolomic data as well as Eugène Maurey for the critical statistical analysis of morphophysiological data.

## 8. Author contributions

GG, CV and AB, designed and assembled the PP genetic construction. AB, MR, CV, AF and GG participated to transgenic lines regeneration. AB, AF and MR cultivated and harvested the tomato plants. AB, FH and RL generated and analyzed the transcriptomic data. AB, JG and RL generated and analyzed the metabolomic data. AB, LH, JR and WB analyzed the lignin composition. AB, CR and RL generated and analyzed the morphophysiological data. AB, RL and AH conceptualized the project and wrote the article. WB, AH and RL supervised the project. All co-authors reviewed the article.

## 11. Supplemental data

**Figure S1:**
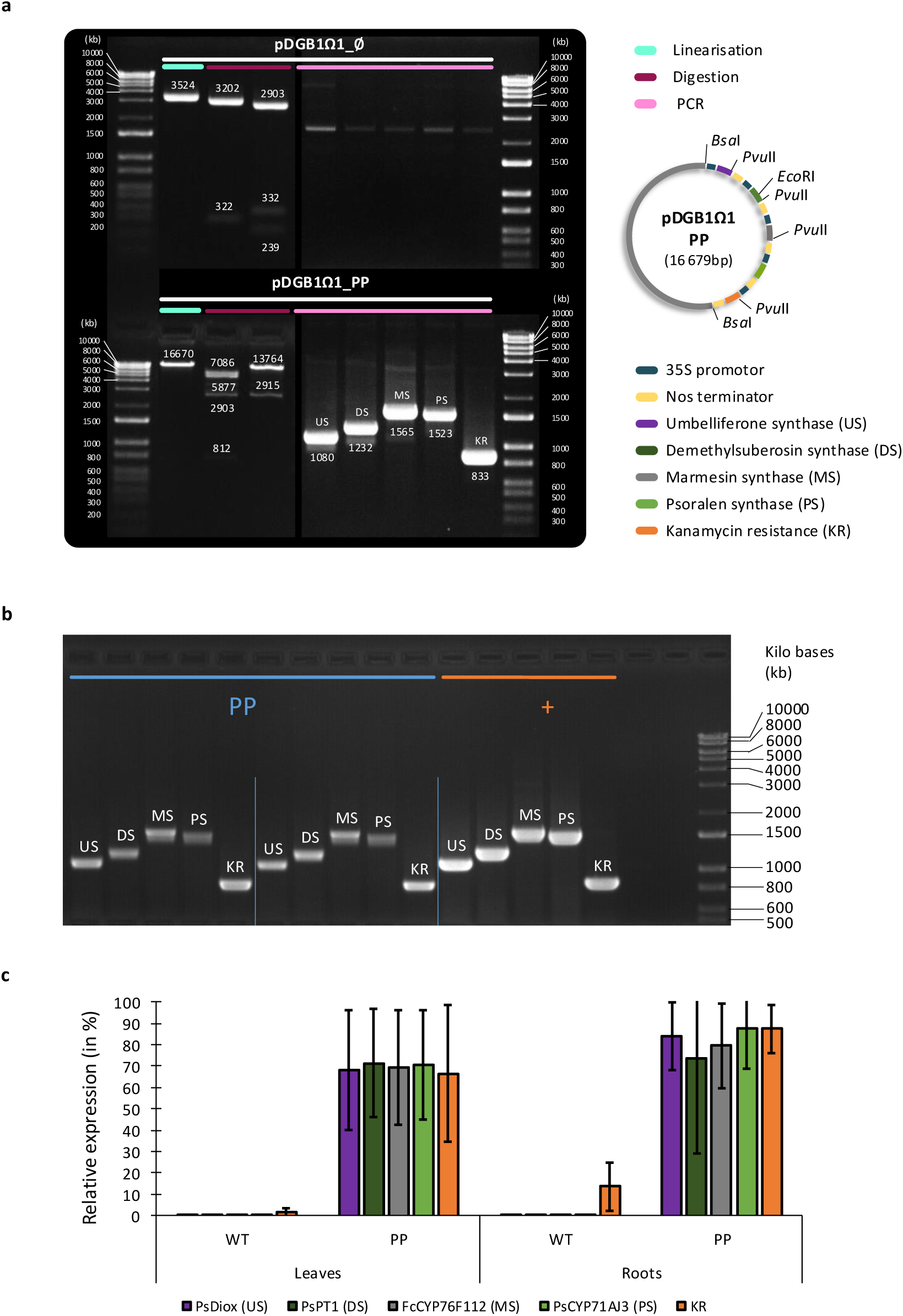
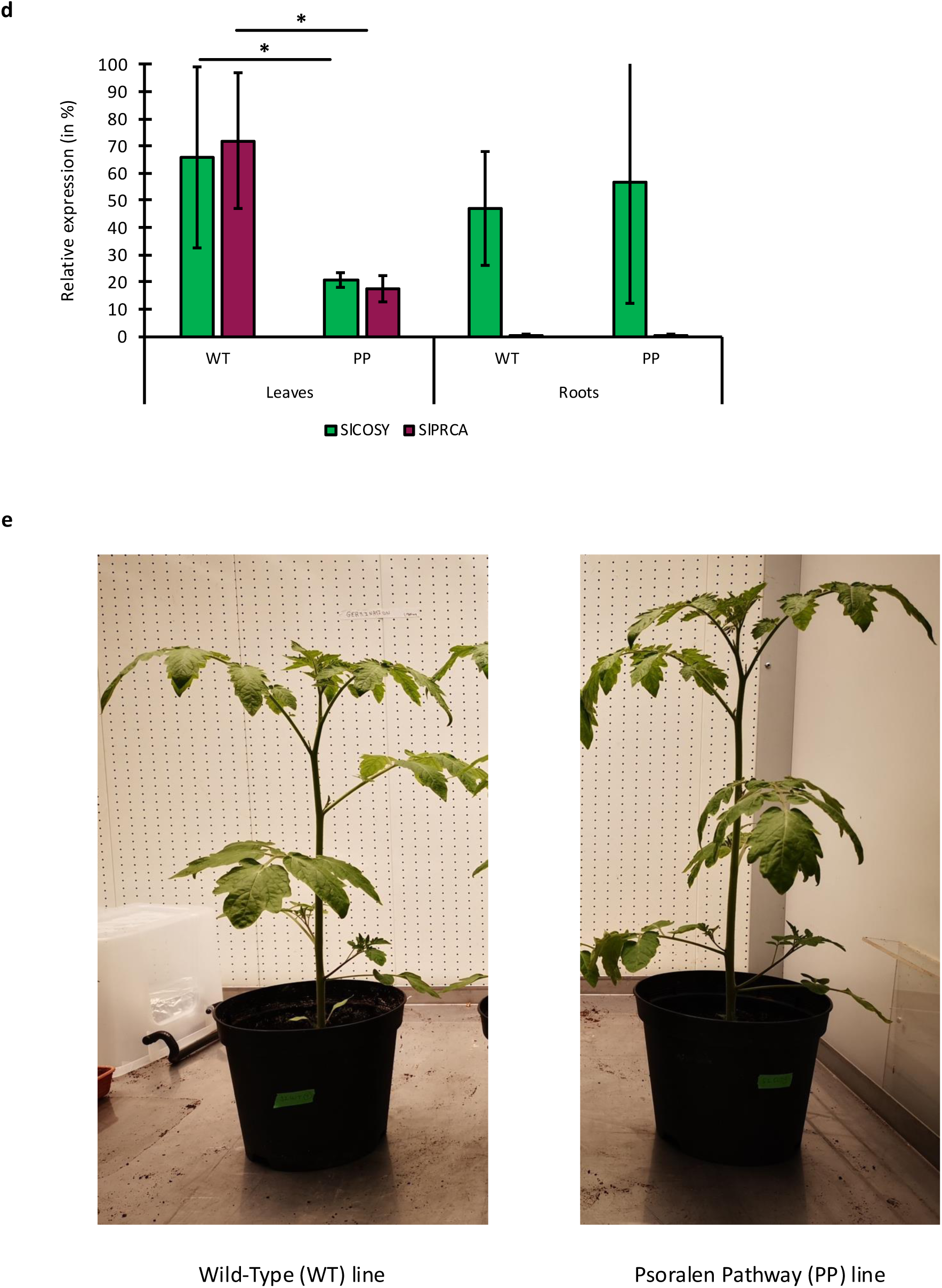
Establishment of the Psoralen Pathway (PP) line. (**a**) Electrophoresis gel verification after linearization, restriction, and PCR of the pDGB1Ω1_PP construct and the empty vector pDGB1Ω1_∅ as a negative control. (**b**) Gel electrophoresis of PCR fragments verification of two T1 transformed lines using genomic DNA for the 5 genes of the PP construct. Positive control (+) is a PCR amplification of the pDGB1Ω1_PP plasmid under the same conditions. (**c**) Expression validation of the five PP transgenes using cDNA from roots and leaves of the WT and PP lines for each conditions in triplicate. (**d**) Expression validation of two genes (*Sl*RCA: Solyc10g086580; *Sl*COSY: Solyc06g082860) significantly down– and upregulated in the PP line according to transcriptomic data, respectively. Actin and GAPDH were used as housekeeping reference genes. (**e**) Photographs of representative tomato plants (*Solanum lycopersicum* var. Wva106, Solanaceae) from the Wild-Type (left) and PP (right) lines, 28 days post-sowing under controlled, non-limiting growth conditions.

**Figure S2:**
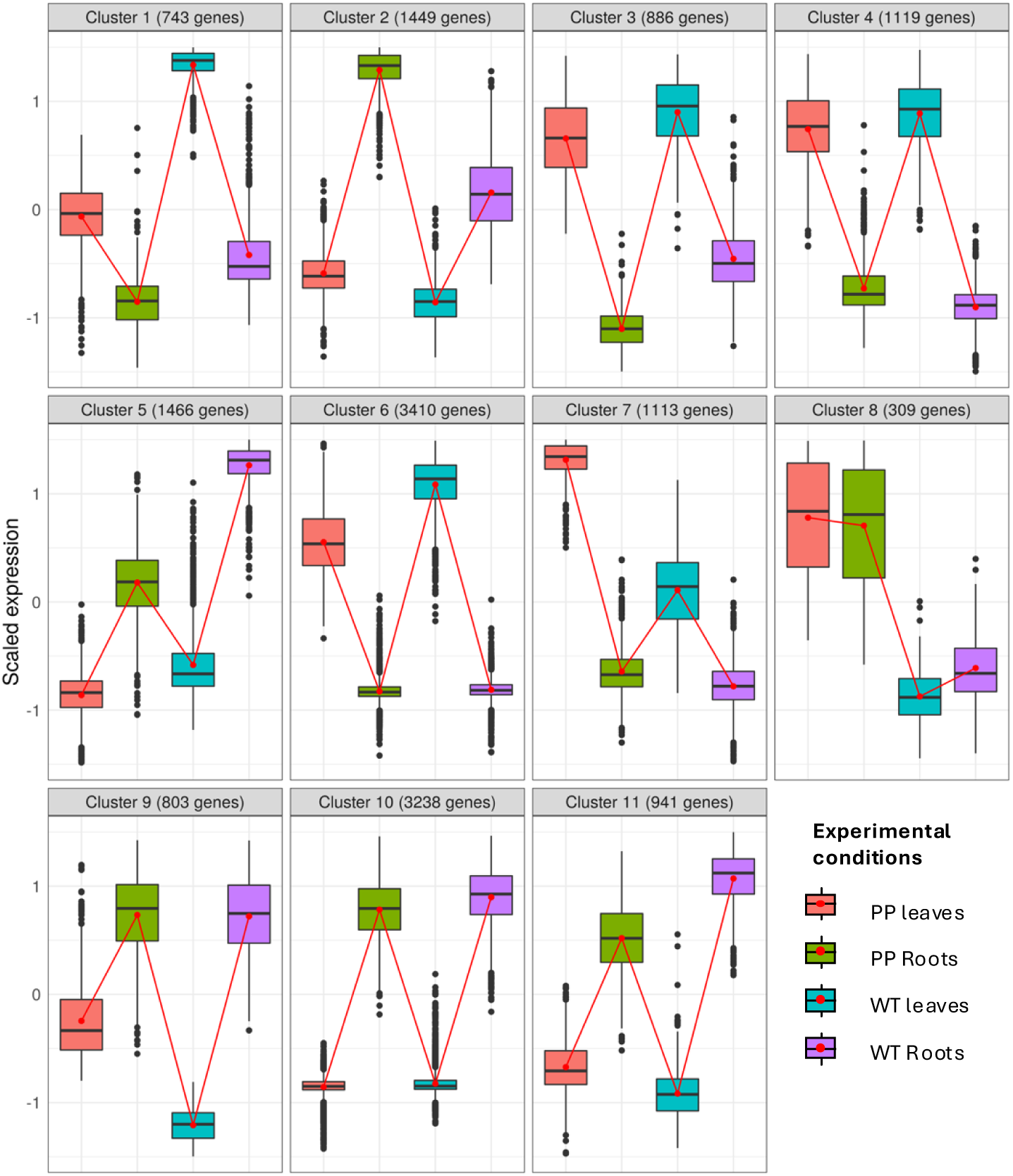
Clustering analysis of the specific gene regulation between the Psoralen Pathway (PP) line and the Wild-Type (WT) line in roots (n=3) and leaves (n=3). Legend: UT: The PP line; WT: WT line; F: leaves; R: Roots.

**Figure S3:**
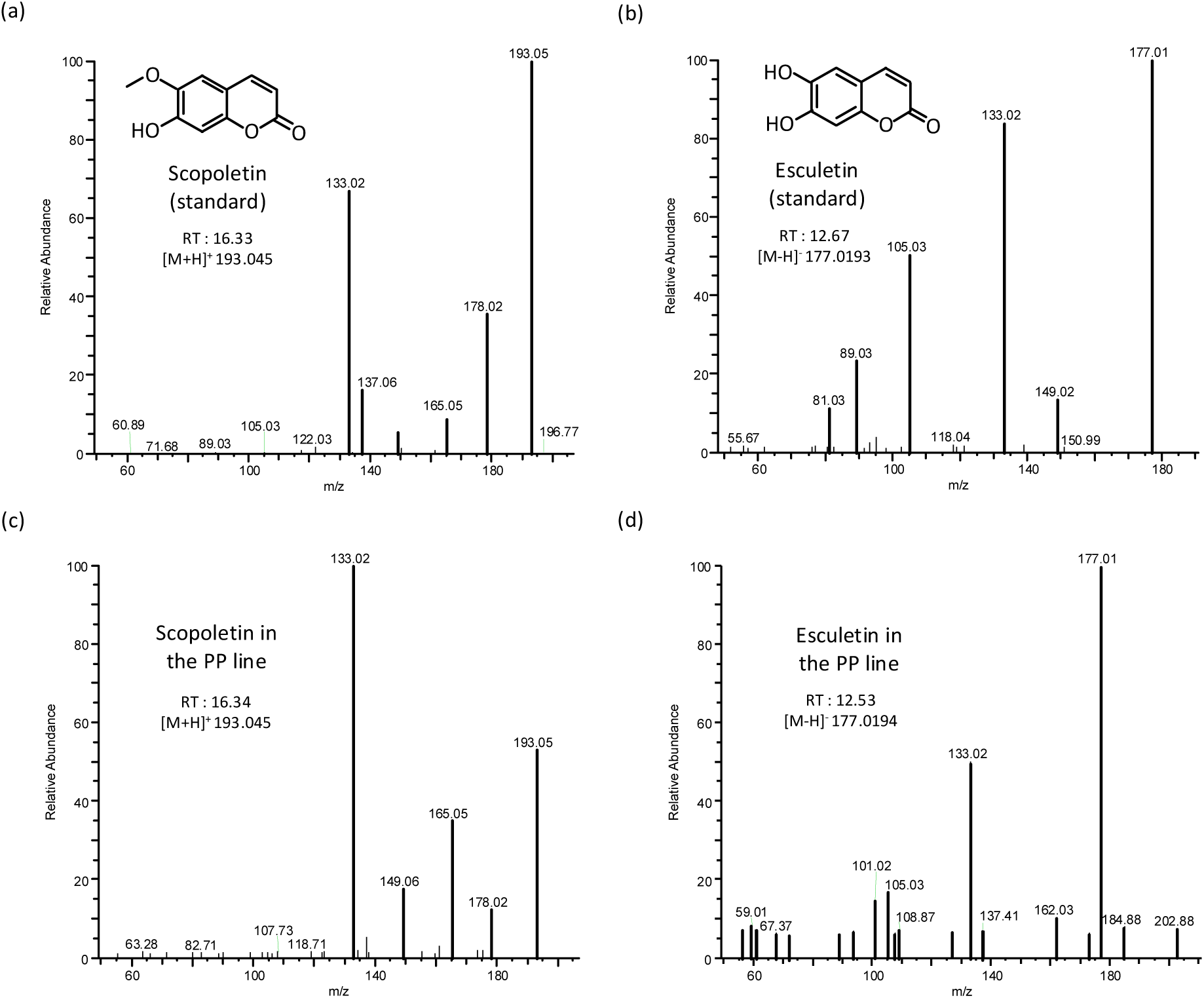
Formal identification of scopoletin and esculetin in the PP line. Tandem mass spectrometry fragmentation pattern of standards of (**a**) scopoletin and (**b**) esculetin and (**c**-**d**) their respective forms identified in the PP line. Analysis was performed after negative (for esculetin) or positive (for scopoletin) electrospray ionization (ESI) and Targeted Single Ion Monitoring (SIM) mode between 50 and 210 m/z. The Higher Energy Collisional Dissociation (HCD) fragmentation mode was investigated.

